# A high-quality draft genome assembly of the Neotropical butterfly, *Batesia hypochlora* (Nymphalidae: Biblidinae)

**DOI:** 10.1101/2025.05.17.654659

**Authors:** Tan Pham Nhat, Anne Duplouy, Joseph See, Lucy S. Knowles, Edgar Marquina, Geoffrey Gallice, Freerk Molleman, Vicencio Oostra

## Abstract

We report a long-read high-coverage reference genome assembly of the Neotropical butterfly, *Batesia hypochlora* (Nymphalidae: Biblidinae). This represents the first reference genome in the Biblidinae subfamily, a clade subject to ongoing studies on seasonal and climate adaptation in the Amazon. We assembled the genome from PacBio HiFi long reads (66X coverage), polished it with Illumina short reads (15X coverage), and annotated it using PacBio IsoSeq RNA data. We observed 15 chromosome-sized scaffolds varying in length from 13.2 Mbp to 37.6 Mbp (median 24.3 Mbp), combining a total genome size of 395.788 Mbp. This assembly is highly contiguous (contig N50 of 25.14 Mbp) and complete (BUSCO completeness score of 98.6% and 0.2% duplication rate). Repeat annotation revealed that the genome consists of about one-third transposable elements. Gene prediction using RNAseq evidence uncovered 19,395 genes, of which 17,400 were assigned to 2,883 orthogroups when including genomes of the fruitfly, silk moth, and three other Nymphalid butterfly species. The high sequencing depth also allowed us to assemble the genomes of the mitochondria and the common endosymbiotic bacterium *Wolbachia*. The mitochondrial genome was fully assembled (15,540 bp in size) with all expected genes annotated. The *Wolbachia* genome was fragmented, and we determined that it belongs to the B-supergroup. The high-quality assembly of *B. hypochlora* can represent the subfamily in further comparative analysis of evolution and provide a key resource for ongoing work to explore reproductive biology and adaptations to seasonality in Amazonian butterflies.

## Background

*De novo* genome assemblies are a key resource for genetic adaptation, evolution, and co-evolution studies. They provide a precise genome map (Sohn & Nam, 2016), for which annotations identify the position of coding genes and other DNA features in the genome, and functional analysis identifies the putative function of these genes (Ranz et al., 2021). *De novo* genome assemblies also provide the backbone of population genomics studies, phylogeography, and demography (Mérot et al., 2020; Yan et al., 2021). Through comparative genomic analyses, they provide deep insights into the macroevolution of organisms. Moreover, new research directions have emerged with the recent shift from fragmented short-read assemblies toward highly contiguous chromosome-scale assemblies based on long reads. This includes quantifying the role of recombination and structural variation for adaptation, the importance of alternative splicing in environmental responses, the identity and nature of cis-regulatory elements underlying ecological adaptations, and comparative phylogenetic analyses, including gene activities in regulatory evolution (van Dijk et al., 2023). The genome of symbionts can be sequenced, assembled, and analyzed along with their hosts. Recently, sequences from such whole genome projects have also been used for metagenomic analyses and identifying non-target organisms like endosymbionts. This brings new insight into the co-evolution between hosts and a wide diversity of parasitic or mutualistic microscopic organisms. In insects, for example, symbiotic bacteria like *Wolbachia* can play an important role in their hosts (Ahmed et al., 2016) and ecology (Russell et al., 2009). When not discarded or ignored, these non-target assemblies can further enrich our understanding of both vertical and horizontal transmission of *Wolbachia* (Ahmed et al., 2016; Lohman et al., 2008; Miyata et al., 2024).

Due to their vast diversity, conservation genomics studies of insects are facing significant challenges, including the scarcity of complete reference genomes. To date, the genomes of 2,596 insect species have been assembled up to the scaffold level and are publicly available in the NCBI database (January 9, 2025). Although Lepidoptera are widely used as models in evolutionary and ecological studies or as biodiversity indicators, only 963 (0.6%) Lepidoptera species have been assembled to the scaffold level, including just 304 (1.8%) of an estimated 20,000 butterfly species globally (Shirey et al., 2022). Thus, most clades remain unrepresented or underrepresented (Bortoluzzi et al., 2023). Of the butterfly genome assemblies available, most come from species in North America or Europe. For instance, although the Neotropics are the world’s most important center of butterfly diversity, with at least 7,000 species (Garwood & Jaramillo, 2022), fewer than 100 butterflies from this region have had their genomes assembled. Therefore, high-quality reference genomes are missing for tropical biodiversity hotspots, which limits further studies across various fields.

One of the clades that lacks a genome assembly is the subfamily Biblidinae. This clade encompasses more than 300 species that arose about 37 Mya and are distributed in South and Central America, with some groups in the Old World tropics (Kawahara et al., 2023). These butterflies are colorful and can be of local economic importance as crop pests (Dias et al., 2012; Dias et al., 2014; Francesconi et al., 2013; Pacheco Gómez et al., 2022). Furthermore, several species are commercially traded to be displayed in live butterfly exhibits or mounted forms (Wang et al., 2023). They have also been the object of studies on wing pattern evolution (Garzón-Orduña et al., 2024), sound production (Yack et al., 2000), and life history (Vasquez J. et al., 2012). More recently, our lab has started using multiple species in this subfamily in a research programme on life history and behavior in the context of adaptation to seasonality and climate change. Therefore, a reference genome assembly for this clade is very timely.

We chose one of the most iconic Amazonian butterfly species to represent Biblidinae, *Batesia hypochlora* C. Felder & R. Felder, 1862 (NCBI id: 127305) is distributed across the western lowland Amazonian rainforest, from central Colombia through southeastern Peru and western Brazil (DeVries et al., 1999; Schoch et al., 2020). It is associated with both intact forest interior and disturbed or edge habitats (Ramos-Artunduaga et al., 2021; GG pers. obs.). The butterfly is likely aposematic as it has a highly conspicuous color pattern and slow flight (DeVries et al., 1999). Moreover, the caterpillars feed on *Caryodendron orinocense*, an evergreen tree belonging to the milkweed (Euphorbiaceae) family (known for its toxic chemicals that, e.g., aposematic monarch butterflies sequester). Adults feed on the juices of rotting fruits (DeVries et al., 1999). However, all other aspects of the species’ biology remain unexplored throughout its range, including reproductive biology, interactions with parasites, and any other interspecific interactions beyond the single known host species.

Here, we provide a reference genome for *B. hypochlora* and its *Wolbachia* endosymbiont. We combine long and short DNA reads to provide a high-quality *de novo* reference genome for *B. hypochlora* and use RNA evidence to complete gene annotations. Our assembly includes chromosome-scale nuclear scaffolds with gene and repeat element annotation, as well as functional annotation. We also provide a complete and annotated mitochondrial genome and a draft assembly of the associated bacterial symbiont *Wolbachia*. Finally, we compared the newly assembled genome with the high-quality reference genomes of the subfamily Nymphalinae (Nymphalidae), including *Aglais io* and *Melitaea cinxia*.

## Material and methods

### Sampling, library extraction, and sequencing

The specimens used in this study were collected at the Manu Learning Centre, a field station located along the Upper Madre de Dios River at an elevation of ca. 450 m a.s.l, where the lower foothills of the Andes meet the lowland the Amazon basin in southeastern Peru (Madre de Dios region; 12°47’18.9”S 71°23’29.1”W). We collected two adults of *B. hypochlora* using fruit- and fish-baited traps suspended at various heights along trails in the primary forest at the site in November 2021. We sacrificed the first specimen (PE-2021-005-R, unknown sex) and stored it in RNAlater, and the second individual (PE-2021-004-E, male) in 100% ethanol. We kept both samples at approximately 4 °C, then transported them at ambient temperature and stored them at -20 °C upon arrival in the laboratory

We used the thorax of the first individual (PE-2021-005-R) for both PacBio DNA (HiFi) and RNA (Isoseq) sequencing (at the NERC Environmental Omics Facilities (NEOF)). We extracted HMW DNA using the Nucleobond HMW kit (Machery-Nagel), with the following modifications: lysis buffer, proteinase K, RNase A, and doubled binding buffer volumes compared to the kit’s protocol. We quantified the HMW DNA using a Qubit fluorometer and measured purity using a Nanodrop (both Thermo Fisher). To assess the integrity, we used Femto Pusle (Agilent), confirming that most of the DNA was > 50 kbp in length. RNA was extracted using the QIAGEN RNeasy Mini Kit. The tissue was initially homogenized using a TissueLyser II (QIAGEN), and the rest of the process was completed per kit instructions. Then, HiFi and Isoseq libraries were prepared and sequenced over 2 SMRT cells (HiFi library) and 1 SMRT cell (Isoseq library)at NEOF Centre for Genomic Research. Combining both SMRT cell data, we obtained 2,337,639 HiFi reads with a median length of 11.4 kbp and 78% of reads with Q30 or more. For the Isoseq reads, we obtained 2,336,880 reads with a median length of 1,882 bp. We used the second individual (PE-2021-004-E) for Illumina short-read DNA sequencing. Half of one thorax was removed from ethanol, dried overnight, and then homogenised using a Tissue Lyser in lysis buffer, and then DNA was isolated using a Qiagen DNeasy Blood & Tissue kit following the manufacturer’s recommendations, including 3-hour Proteinase K incubation. Short insert size (ca. 350 bp) Illumina sequencing libraries were prepared (NEBNext Ultra II FS Kit with ½ volume reactions). The sample was sequenced with other samples on a Novaseq S4 lane (150 bp PE, yielding 63.5M raw reads (30.1M reads after filtering) at NEOF Centre for Genomic Research

### Quality control and pre-assembly estimates

To check the quality for both HiFi long reads and Illumina short reads, we used FastQC v0.11.9 (Andrews, 2010). According to the results from the given FastQC, only paired-end Illumina short reads have adapters, so we used trimmomatic v0.36 (LEADING: 3 TRAILING: 3 SLIDINGWINDOW: 4: 20 MINLEN: 36) to remove adapters and low-quality data (Bolger et al., 2014; Tilak et al., 2018), and then the trimmed reads were rechecked. We performed a k-mer analysis on HiFi reads to estimate genome size, heterozygosity, repetitiveness, and sequencing coverage. We used Jellyfish v2.2.10 (Marçais & Kingsford, 2011) to calculate 31-mer normalized coverage, then visualized and estimated parameters using Genomescope v2 (ploidy p=2 and kmer=31; at qb.cshl.edu/genomescope/genomescope2.0/) (Vurture et al., 2017).

### Nuclear genome assembly

We used HiFi long reads for initial assembly using hifiasm v0.19.5-r587 with default parameters (Cheng et al., 2021), yielding genome version 0.1. We then polished genome version 0.1 with both PacBio long reads and Illumina short reads. First, we aligned the PacBio long reads using minimap2 (Li, 2018) and trimmed Illumina short reads using bwa (Li, 2013) to the assembly version 0.1. We then used these alignments to polish the version 0.1 using Pilon v1.24 (--fix gaps,local,breaks) (Walker et al., 2014), resulting in version 0.2. We identified and removed xenobiotic contamination (*Wolbachia*, see Results) using Blobtoolkit v4.2.1 (Challis et al., 2020), resulting in assembly v.0.3.

We calculated contiguity statistics and completeness for each assembly version. The module stats of bbtool v39.01 generated basic statistics, including scaffold count, N50, L50, and gap percent (Bushnell, 2015), while completeness was calculated using both Busco v5.5 (Manni et al., 2021) and Compleasm v0.2.2 (Huang & Li, 2023) against the lepidoptera_odb10 database.

### Mitochondrial genome assembly

We assembled the mitochondrial genome using MitoHiFi v3.2 from assembly v.0.3 (Uliano-Silva et al., 2023). At the beginning of the mitochondrial assembly, we used

findMitoReference.py to determine a close relative reference mitochondrial genome to *B. hypochlora*, and it downloaded a completed mitochondrial genome of *Hamadryas epinome* (NC_025551.1) (Nymphalidae: Biblidinae) as the most closely related species for which a mitochondrial genome is available (Cally et al., 2016). We finally ran mitohifi.py for mitochondrial annotation and reported the mitochondrial genome separately from the final nuclear genome. The remaining nuclear genome (without the mitochondrial genome) was named assembly version 0.4.

### Repeat, Gene, and Functional annotations

We identified repeats *de novo* in assembly version 0.4 for using the database of RepBaseRepeatMaskerEdition-20181026 with RepeatModeler v2.0.5 (Bao et al., 2015; Flynn et al., 2020; Smit et al., 2024). We then split the library into known (successfully classified) elements and unknown elements (remaining unclassified or unknown) using seqkit (Shen et al., 2016). Then, we used RepeatMasker v4.1.5 to mask repetitive elements from the Repbase and Insecta repeat libraries using repclassifier v1.1 (Card, 2022; Smit et al., 2024). Unique transposable element (TE) families were grouped into eight different TE classes, including “DNA Transposons”, “Helitrons”, “LINEs” (Long interspersed nuclear elements), “LTR Retrotransposons” (Long terminal repeats), “Low Complexity”, “SINEs” (Short interspersed nuclear elements), “Simple Repeat”, and “Unknown”. We present the soft-masked genome as the final assembly (v.1.0).

We combined the soft-masked genome version 1.0 with RNA Isoseq data to predict gene models. We followed the Isoseq workflow to generate both high-quality (predicted accuracy ≥ 0.99) and low-quality (predicted accuracy < 0.99) reads, removed primers and barcodes from the raw of Isoseq reads by using lima, and then refined (by removing polyA tail) and clustered (Pacific Biosciences, 2019). Then, we aligned only high-quality RNA reads to genome version 1.0 using minimap2 (Li, 2018). We then used Braker v3.08 to predict proteins using RNA alignment bam file, soft-masked genome assembly (version 1.0), and training with the protein sequence of Arthropoda from OrthoDB v.11 (Brůna et al., 2024; Gabriel et al., 2023; Kuznetsov et al., 2022).

We used two methods to annotate the function of the protein-coding genes. The first method involved identifying *B. hypochlora* orthogroups in 5 other insects (*Drosophila melanogaster*, *Bombyx mori*, *Danaus plexippus*, *Heliconius melpomene*, *Melitaea cinxia*) using Orthofinder (Emms & Kelly, 2019). Subsequently, we downloaded the UniProtKB database for these five insects. This yielded protein-to-GO mappings for all *B. hypochlora* proteins with an annotated ortholog in at least 1 species. The second method was to identify Gene Ontology (GO) in the genome of *Batesia* (version 1.0) by mapping the gene sequence to the precompiled database for insects. We downloaded the database for “Insecta” taxa on eggNOG DB v5.0.2 using create_dbs.py (Huerta-Cepas et al., 2017) and annotated the *Batesia* protein sequences using eggNOG-mapper v2 (Cantalapiedra et al., 2021). Then, we used an online version of GOTermMapper (https://go.princeton.edu/cgi-bin/GOTermMapper) to map the unique GO terms to GO slim based on the Ontology aspects (Boyle et al., 2004; Gene Ontology Consortium, 2004). Finally, we integrated the results of both methods to obtain the gene function.

### Microbial symbiont detection and assembly

We used the Blobtoolkit (Challis et al., 2020) to analyze possible non-host genomic material, including sequences from common endosymbiotic bacteria, such as *Wolbachia* and *Spiroplasma* (Duplouy & Hornett, 2018). We identified 27 *Wolbachia* contigs from the host nuclear and mitochondrial assemblies (ie. version 0.3, see Results). To further characterize these at the strain level, we isolated and screened them for the presence of the *wsp* gene and the five Multi Locus Strain Typing (MLST) genes using the blast function in Geneious Prime® 2025.0.3 (https://www.geneious.com). These six loci are commonly used to phylogenetically assign *Wolbachia* strains to their respective taxonomic supergroup (Baldo et al., 2006). The five MLST genes (ie. *ftsz, fbpa, coxA, gatB,* and *hcpa*) were all identified and compared to orthologous genes from reference genomes of different *Wolbachia* supergroups in gene-specific phylogenetic analyses following the protocol described in (Twort et al., 2022). In brief, a reference set for each gene was obtained from GenBank with representative strains of the *Wolbachia* A-, B-, F-, and D-supergroups (Twort et al., 2022). Individual gene alignments were produced using the pairwise alignment with the default options in Geneious Prime. Alignments were manually screened to check and correct any errors. Phylogenetic reconstructions and tree visualizations of *Wolbachia* supergroups were carried out for each locus independently using the Tree function with default options in Geneious Prime.

We similarly screened for the presence of the *wmk* and *Oscar* genes (Katsuma et al., 2022; Perlmutter et al., 2019) and of both the *cifA* and *cifB* genes (Beckmann et al., 2017; LePage et al., 2017) in our *Wolbachia* contigs, using the BLAST function with default settings in Geneious Prime. We used the WolWO-mediated killing-like protein (*wmk*) gene sequence from the *Wolbachia* strain *w*CauB (#MK955149.1), the amino acid sequence and domain structure of the Oscar protein given by Katsuma et al (2020), and the cytoplasmic incompatibility factor A and B protein genes (#MG807657, #MG807658, OP947615, and #MH544806) from *Wolbachia* strain *w*Pip. The detection of the *wmk* and *Oscar* genes, the candidate genes for the expression of the male-killing phenotype in *Wolbachia* (Katsuma et al., 2022; Perlmutter et al., 2019) could suggest that the *Wolbachia* strain, which infected *B. hypochlora* (labeled: *w*Bhyp), induces the death of the male progeny of its host (Duplouy et al., 2013; Perlmutter et al., 2019). Similarly, the detection of the *cifA* and *cifB* genes, which code for the expression of cytoplasmic incompatibility (CI) between individuals of incompatible infection status (Beckmann et al., 2017; LePage et al., 2017), could suggest that *w*Bhyp can induce CI in *B. hypochlora*.

### Large-scale genome rearrangement analysis

The soft-masked genome of *B. hypochlora* (version 1.0) was compared to a published high-quality reference genome to determine any large-scale genome rearrangements in the genome. As no other genome of the Biblidinae subfamily is available, we used more distantly related species with a published chromosome-level genome assembly in the subfamily Nymphalinae. We selected *Aglais io* (Nymphalini, Nymphalinae, NCBI ID: 171585) and *Melitaea cinxia* (Melitaeini, Nymphalinae, NCBI ID: 113334). These species have a divergence time of ca. 57 Mya, and for both, the sex chromosomes have been identified (Challis et al., 2023; Kawahara et al., 2023). We downloaded the reference genomes from the NCBI database and only retained (for both query and references) scaffolds or chromosomes longer than 1 Mbp. We used nucmer in MUMmer v4.0.0rc1 for alignment with a set of a minimum length of cluster of matches (c) is 100, and a minimum length of a single exact match (l) is 500bp (Marçais et al., 2018). The resulting nucmer alignment was visualized with the “circlize” or “ggplot2” packages in R 4.2.2.

## Results

### Nuclear and mitochondrial genome assemblies

Based on the k-mer analysis of the PacBio long read, the nuclear genome of *B hypochlora* has an estimated haploid length of 370 Mbp and a low heterozygosity of 0.43% (Supplement 1, Figure S1). The total genome size in the assembly version 0.1 is 397.812 Mbp in 143 contigs, with the longest contig being 37.56 Mbp and a contig N50 value for the genome assembly of 25.29 Mbp (Supplement 1, Table 2). During the polishing process, we corrected the position of 863,381 bp in 321 locations, including fixing breaks (deletion of 378,316 bp and insertion of 71,092 bp) and opening gaps (deletion of 456,931 bp and insertion of 73,739 bp). This resulted in an assembly size of 397.124 Mbp (version 0.2). The xenobiotic analysis identified 27 non-Lepidoptera scaffolds representing 1.34 Mbp of the genome. These were all version 0.2 assigned to Pseudomonadota bacteria (Supplement 1, Figure 2), which turned out to be *Wolbachia* (see detail below). After removing the *Wolbachia* contigs, the final genome size of *B. hypochlora* is 395.788 Mbp, with 99.19% of the main genome in 16 scaffolds >= 1 Mbp (genome version 0.3, Figure 1).

**Figure 1.**
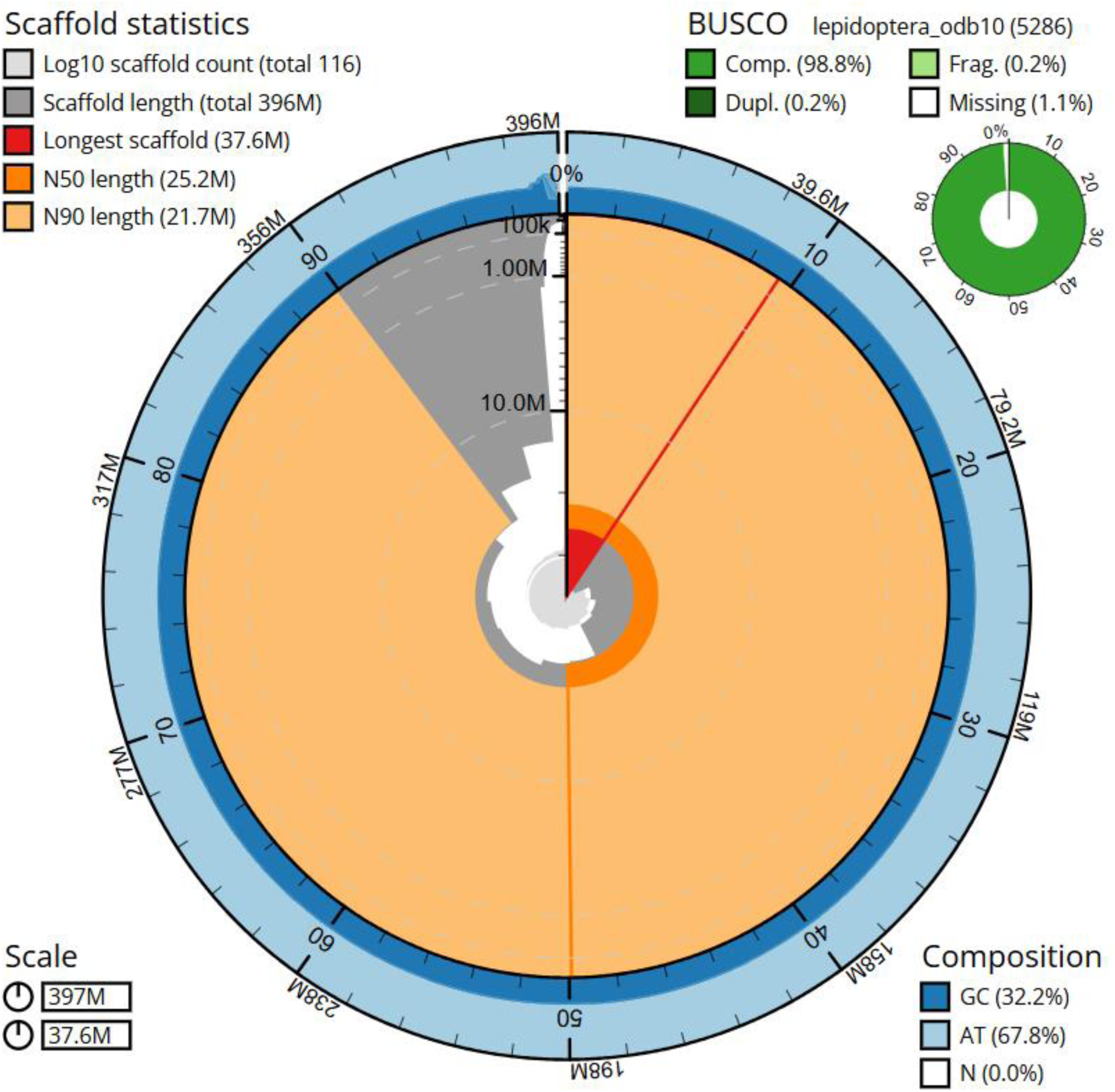
Snail plot summary of assembly statistics for genome assembly version 0.3. (after removing *Wolbachia*, Supplement 1, table 3), generated on the Galaxy Server. The main plot is divided into 1,000 size-ordered bins around the circumference with each bin representing 0.1% of the 397,123,821 bp assembly. The distribution of sequence lengths is shown in dark grey with the plot radius scaled to the longest sequence present in the assembly (37,552,201 bp, shown in red). Orange and pale-orange arcs show the N50 and N90 sequence lengths (25,221,893 and 21,698,544 bp), respectively. The pale grey spiral shows the cumulative sequence count on a log scale with white scale lines showing successive orders of magnitude. The blue and pale-blue area around the outside of the plot shows the distribution of GC, AT, and N percentages in the same bins as the inner plot. A summary of complete, fragmented, duplicated, and missing BUSCO genes in the lepidoptera_odb10 set is shown in the top right. The assembly has been filtered to exclude sequences with family matches Anaplasmataceae (genus matches *Wolbachia*)

**Figure 2.**
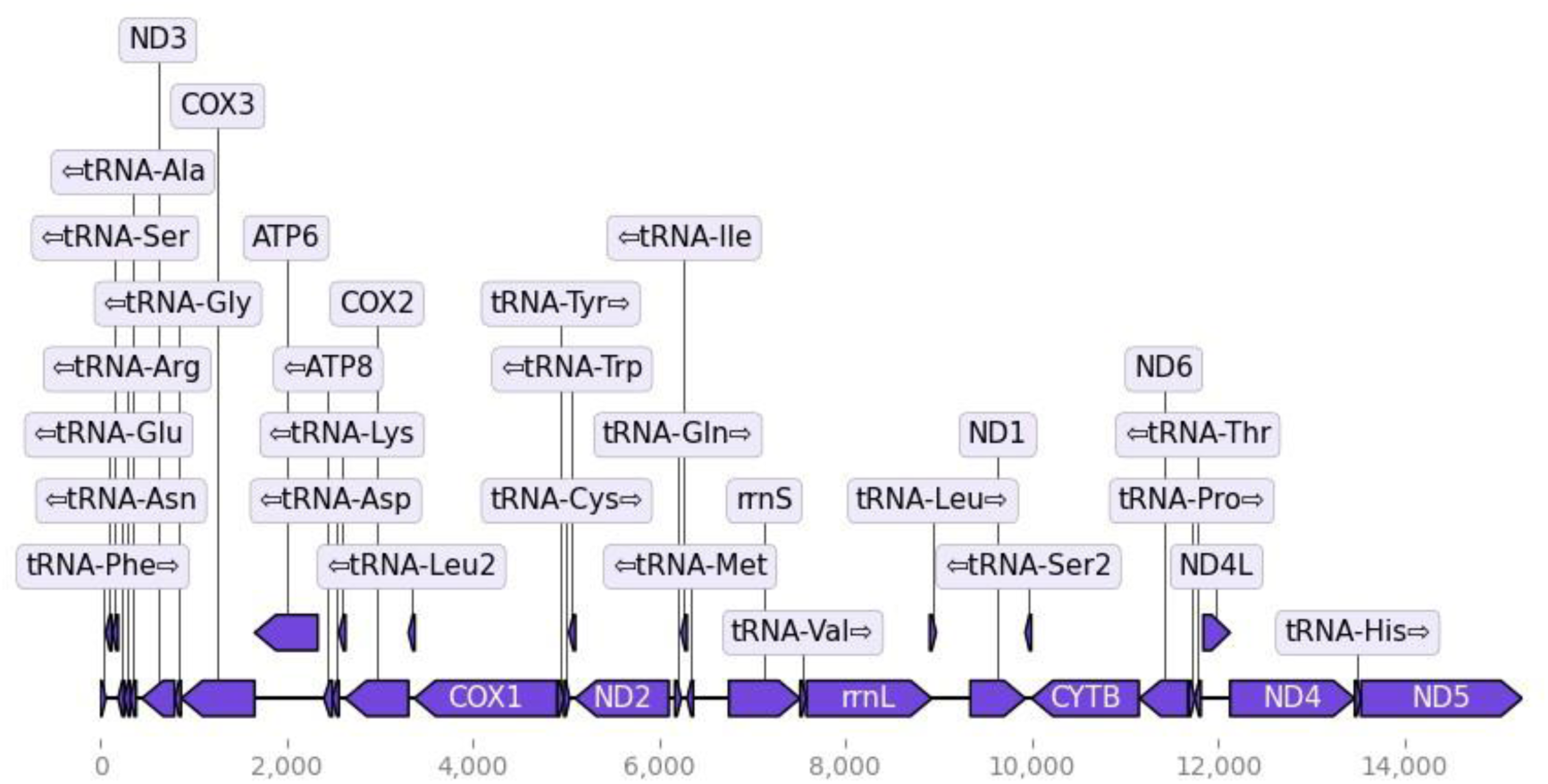
Annotation of mitochondrial genome of *Batesia hypochlora*. The horizontal axis shows the position across the mitochondrial genome (total length 15,521 bp), with the location of 13 coding genes, 22 tRNAs, and 02 rRNAs (rrnS & rrnL) indicated in purple and the transcription direction of each gene indicated with an arrow. Plot generated using MitoHiFi (Uliano-Silva et al., 2023)

The complete mitochondrial genome of *B. hypochlora* consists of 15,540 bp with an order of 37 genes, including 13 protein-coding genes, 22 transfer RNAs, and 2 rRNAs (Figure 2). The majority strand (J-strand) has 23 genes (9 PCGs and 14 tRNAs), while the minority strand has 14 genes (4 PCGs, 8 tRNAs, and 2 rRNAs; Supplement 1, Table 6).

We separated the mitochondrial genome from the nuclear genome version 0.3 so that the total genome remained at 395.788 Mbp in genome version 0.4 (Supplement 1, Table 2). The assessment of completeness using BUSCO and Compleasm from genome version 0.1 to genome version 0.4 was higher than 98.7%, with a duplication rate of 0.2% (Supplement 1, Table 4). After repeat masking (see details below), we got a soft-masked genome as the final genome assembly. The assembly version 1.0 size is 395.788 Mbp (Table 1), including 391.297 Mbp corresponding to 15 chromosome-sized scaffolds (> 2.5 Mbp). This assembly was highly contiguous, with an N90 of 21.698 Mbp (Supplement 1, Table 3).

**Table 1.**
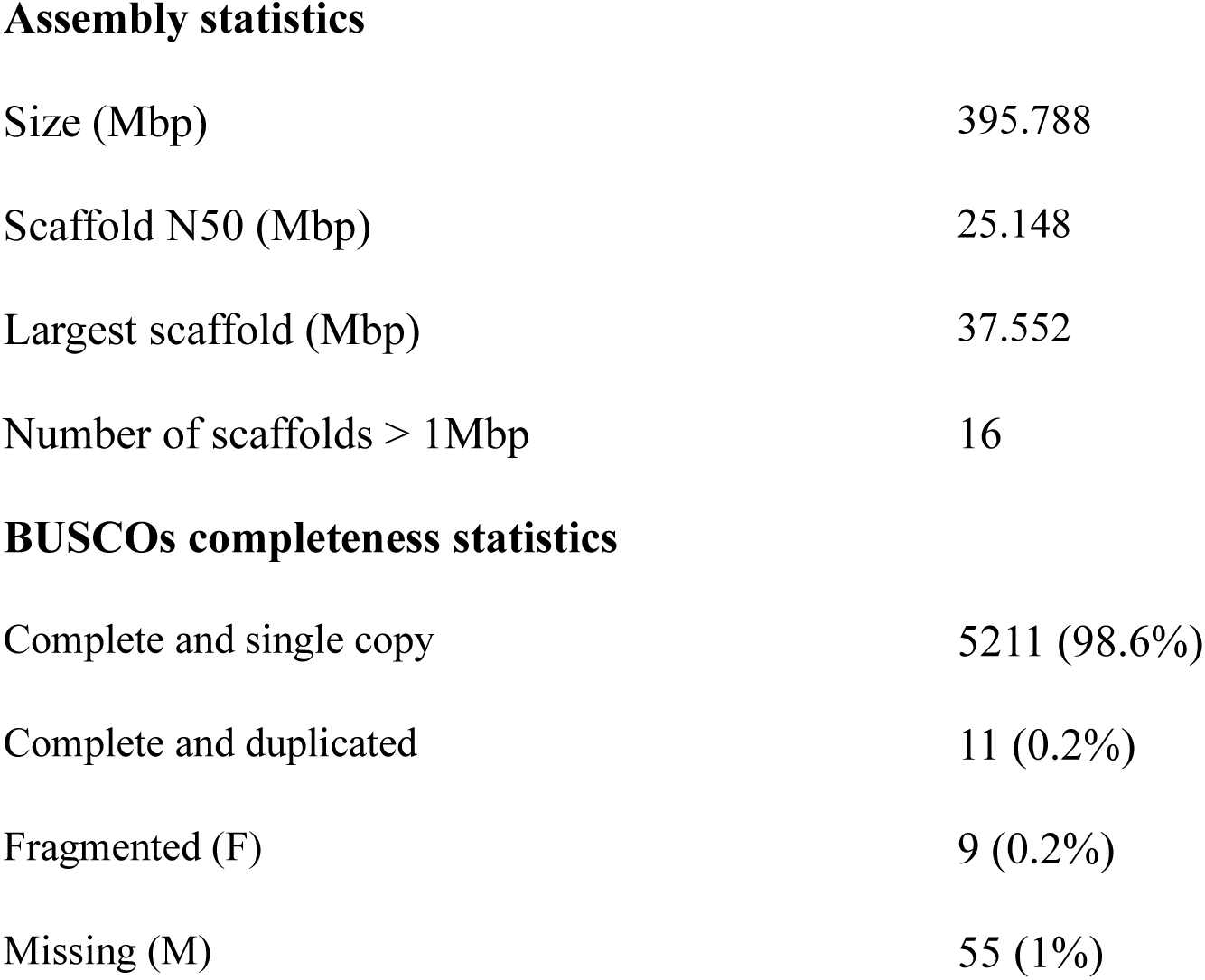
Genome assembly statistics. Evaluation of statistics based on genome version 1.0, and completeness based on genome version 0.4

|  |  |
| --- | --- |
| Size (Mbp) | 395.788 |
| Scaffold N50 (Mbp) | 25.148 |
| Largest scaffold (Mbp) | 37.552 |
| Number of scaffolds > 1Mbp | 16 |
| **BUSCOs completeness statistics** |  |
| Complete and single copy | 5211 (98.6%) |
| Complete and duplicated | 11 (0.2%) |
| Fragmented (F) | 9 (0.2%) |
| Missing (M) | 55 (1%) |

### Microbial symbiont detection and *Wolbachia* assembly

Blast analysis identified 27 contigs as *Wolbachia,* with the largest ca. 100kbp long (average 49.47 kbp), and a combined total length of 1.34 Mbp long. Sequencing depth for reads mapping to *Wolbachia* contigs was 1% and 9% of depth for reads mapping to the host genome for two individuals (Supplement 2). Through phylogenetic analyses of each MLST sequence from the *w*Bhyp *Wolbachia* contigs (Supplement 1, Figure 4), we were able to confidently place *w*Bhyp in the *Wolbachia* B-supergroup (>95% similarity with *w*Pip (B-), ≈ 85% similarity with *w*Mel (A-) and *w*Bm (D-supergroup)). We could not retrieve the *wsp* gene from our partial *w*Bhyp assembly. Similarly, we did not find any evidence of the presence of the *cifA* and *cifB* genes (LePage et al., 2017) nor of the *Oscar* gene (Katsuma et al., 2022) in the *Wolbachia* contigs. However, we were able to retrieve a putative *wmk* gene from *w*Bhyp assembly, suggesting the strain might be able to induce male-killing in its butterfly host *B. hypochlora*.

### Repeat annotation, gene model prediction, and functional annotation

We identified 1135 different types of repeats, classified as 1,042 known and 93 unknown elements, after two rounds of running the repclassifier module, and TEs accounted for 34% of the nuclear genome content in the final *B. hypochlora* nuclear assembly (soft-masked genome, version 1.0). The most abundant repeat class across all TE categories was LINEs (47.85 Mbp, n = 241,677), followed by Simple Repeat (9.81 Mbp, n = 203,837) (Figure 3). DNA transposons account for 2.8 % of the total *B. hypochlora* genome size (11.16 Mbp, n = 34,047).

**Figure 3.**
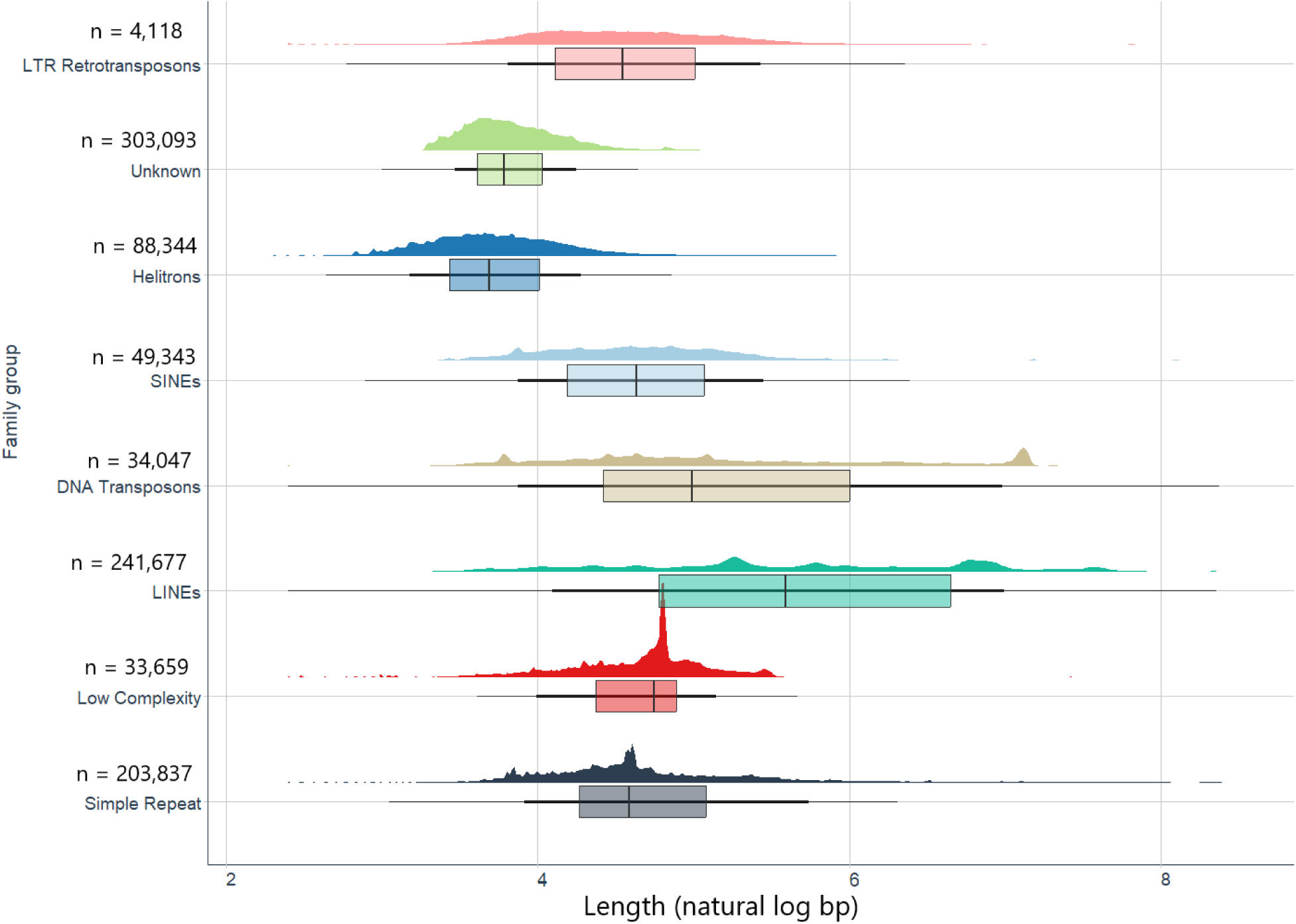
Length distributions of transposable elements in eight TE superfamilies. The horizontal axis is the length (natural logarithm of bp), with the vertical axis showing the frequency. Eight superfamilies are shown from top to bottom in different colors, annotated with their number of elements.

Incorporating RNAseq evidence from the thorax, we identified 21,588 mRNAs of median length 1624.5 bp (min = 6 bp, max = 354,511 bp). We constructed 19,395 gene models (mean length = 4,423.8 bp, median length = 1,461 bp) from the masked genome (version 1.0). This also consisted of 74,395 introns of median length 379 bp (min = 31 bp, max = 211,373 bp) and 95,983 exons of median length 163 bp (min = 1 bp, max = 119,64 bp). The average number of introns and exons per gene was 3.8 and 4.9, respectively.

The eggNOG-mapper detected 18,460 genes with an average of 63.31 GO terms per gene, including 8560 genes containing at least one GO term (mean = 136.5 GO terms, median = 101 GO terms). We mapped 5,646 unique GO terms to GO Slim in three ontology aspects (Supplement 1, Figure 5). Finally, we also used the protein sequence for orthology analysis with five different species (*Drosophila melanogaster*, *Bombyx mori*, *Danaus plexippus*, *Heliconius melpomene*, *Melitaea cinxia*), and OrthoFinder assigned a total of 19,393 genes (81.8 % of all genes in 6 species) to 2,957 orthogroups, with a mean orthogroup size of 6.6. In *Batesia hypochlora* protein sequences, 17,400 genes (80.6%) were assigned to 2,883 orthogroups.

### Large-scale genome rearrangements

Examining chromosomal rearrangements between our *B. hypochlora* genome assembly and *A. io* genome, which shares a last common ancestor ca. 57 Mya (Kawahara et al., 2023), shows strong syntenic relationships across scaffolds (Figure 4). We identified 17,496 alignments between the *B. hypochlora* on 17 scaffolds and all chromosomes of *A.io*. We detected large-scale rearrangements in all 15 chromosome-sized scaffolds (mean length = 610 bp, ca. 2.63% genome size) of the *B. hypochlora* genome between these genomes, except there was no alignment with chromosome 27 (8.7 Mbp) in the *A.io* genome. In general, many scaffolds in the *B. hypochlora* genome appear as fusions of chromosomes compared to *A. io*. For instance, scaffold 1 in the *B. hypochlora* genome appears to have resulted from a fusion of chromosome 7 with an inversion of major portions of chromosomes 20 and 26 of the *A. io* genome. Similarly, scaffold 3 in *B. hypochlora* appears to have resulted from a fusion of chromosome 3 with an inversion of the entire chromosome 13 from the *A. io* genome (Supplement 1, Figure 6). We found that scaffold 14 of the *B. hypochlora* genome was mapped to chromosome Z from the *A. io* genome. We also observed a similar pattern of fusion of chromosomes when we compared *B. hypochlora* and *M. cinxia* genomes, with 14,894 alignments in 15 chromosome-sized (mean length = 605 bp, ca. 2.12% of *B. hypochlora* genome size). The supposed sex chromosome was also mapped to the sex chromosome in M. *cinxia* (Supplement 1, Figures 6 & 7). Moreover, when we mapped the Illumina short reads of our male specimen to the reference genome, the coverage against scaffold 14 (the putative B. hypochlora sex chromosome) was identical to that of the autosomes (Supplement 2), consistent with a ZZ individual. Therefore, we tentatively conclude that scaffold 14 represents chromosome Z of *B. hypochlora*.

**Figure 4.**
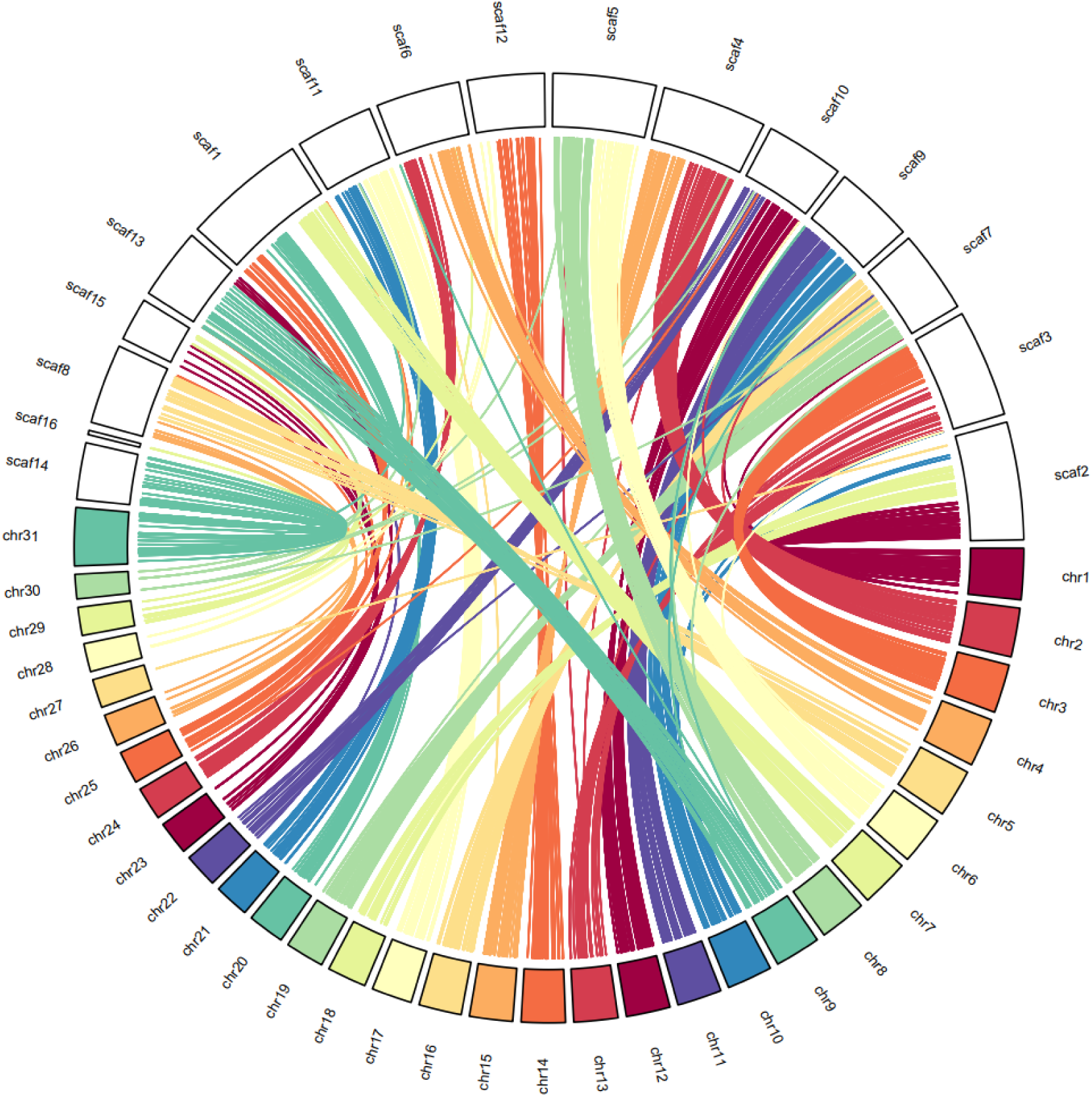
Lare-scale rearrangement and genome evolution compared with the genome of *Aglais io*. Circos plot shows results from the whole-genome alignment using nucmer for contigs > 1 Mbp, with the minimum length of a cluster of matches (c) being 100 bp, and the percentage of identity higher than 80%. Chromosome 31 is the sex chromosome in the *A.io* genome.

## Discussion

We present a high-quality genome assembly for the Neotropical butterfly, *Batesia hypochlora*, the first reference genome in the subfamily Biblidinae. We identified 15 chromosome-sized scaffolds, which is less than half of the reconstructed ancestral chromosome number of 31 for the Biblidinae subfamily and the Nymphalidae as a whole based on cytological counts (Ahola et al., 2014; S. Brown et al., 2007; Wright et al., 2024). We initially considered a technical error, suspecting that our genome assembly may have accidentally fused (fragments from) different chromosomes onto the same scaffold. While we lacked HiC data to exclude this unambiguously, our analysis of coverage variation of PacBio HiFi and Illumina short fragment paired-end reads failed to identify putative incorrect fusion points. Therefore, we conclude that the putative chromosome number of 15 in *Batesia hypochlora* is not a methodological artifact and instead reflects the remarkable biological variation in chromosome number observed across the subfamily Biblidinae as described previously using karyotype analysis (S. Brown et al., 2007). For example, within the same tribe (Ageroniini), representatives of other genera, such as *Panacea* (the most closely related genus) and *Hamadryas,* have the ancestral chromosome number (n=31), but *Ectima* also has about half (n=16). Still, several other tribes show considerable variation, including Epiphilini, which include species with n=7, 10-14, 27-34, and 54, Biblidini (n = 15 and 28-33), and Epicaliini (n = 7-8, 11, 14-16, 21-31). There is even substantial variation within genera and even within species (*Eunica malvina*, n=14 and 31). In contrast, two tribes (Callicorini and Eubagini) exclusively have genera with the ancestral or close to ancestral chromosome number (n=28-31). Thus, overall chromosomal fusions (and to a lesser extent, fissions) are common in Biblidinae, consistent with our finding of n=15 for *B. hypochlora*.

The rate of chromosomal rearrangements in evolutionary time is variable in different groups of organisms (Augustijnen et al., 2024), and this rate is high in butterflies (Coghlan et al., 2005). While we have information about chromosome numbers in some Biblidinae, we do not have another genome in this subfamily to get a glimpse of which parts were rearranged in what fashion. Therefore, we compared the *B. hypochlora* genome with two species from another subfamily of the same family, *A. io* and *M. cinxia*. These chromosome synteny analyses indicated many large-scale rearrangements in the *B. hypochlora* genome, with the joining and inversion of multiple chromosomes. The frequency of chromosomal rearrangements is higher between closely related species, indicating that a chromosomal rearrangement would be a cause for speciation (de Vos et al., 2020; Wright et al., 2024). Chromosomal rearrangements with fusion and fission are common in Nymphalidae, including both cladogenesis and anagenetic events. The anagenetic event rates are higher than cladogenesis event rates in Nymphalidae, except in Ithomiine butterflies (de Vos et al., 2020). Hence, further tests are needed to evaluate the chromosomal rearrangement rate with fusion or fission events during *B. hypochlora* speciation.

With 34%, the observed TE content in *B. hypochlora* fell within the range of variation in Nymphalid butterflies (7% in *Danaus plexippus* to 55% in *Melanargia galathea* (Wright et al., 2024)) and is similar to *M. cinxia* with 42% (Smolander et al., 2022). There is strong evidence that larger genomes tend to have a higher TE content (Talla et al., 2017). Moreover, the percentage of TE in the *Batesia* genome was close to its mean in genomes of species more closely related to *Batesia*, an average of 34.88% with a divergence of 57 Mya (Nymphalinae) and an average of 41.63% with a divergence of 63 Mya (Satyrinae, Heliconiinae, or Limenitidinae) (Wright et al., 2024). The evidence of higher TE contents in a genome indicates higher rates of chromosome fusion (Ahola et al., 2014; Höök et al., 2023). TE frequencies are higher in smaller autosomes than in sex-linked elements (Wright et al., 2024). Further studies are needed in taxa which has a wide range of chromosome numbers as Biblidinae, to demonstrate the relationship between TE content and speciation rate or divergence time.

We obtained 1.34 Mbp of the *Wolbachia* genome during *B. hypochlora* genome assembly. This is similar to the median of *Wolbachia* genome size (ca.1.3 Mbp) in Lepidoptera (Katsuma et al., 2022). The low depth of read mapping observed suggests a low *Wolbachia* titer, which might be expected from thorax tissue and male hosts. As both sequencing approaches used here produced reads for the *Wolbachia* assembly, we can confidently say that the symbiont infected both *B. hypoclora* specimens we sequenced. Butterflies are well-known hosts of *Wolbachia* (Duplouy and Hornett 2018), providing many textbook examples of the role of *Wolbachia* in the ecology and evolutionary histories of these insects (Charlat et al., 2009; Hornett et al., 2008; Salunkhe et al., 2014). Our partial assembly of the *w*Bhyp strain provides evidence that this particular *Wolbachia* strain sits in the *Wolbachia* B-supergroup, which has also been often found in Lepidoptera (Ahmed et al., 2016; Duplouy et al., 2021). Finally, we successfully isolated the *wmk* gene, a putative gene for the expression of male killing in *Wolbachia* (Perlmutter et al., 2019). Although we could not retrieve the *wsp* gene, CI-coding (*cifA* and *cifB*), and the *Oscar* gene in our partial assembly of *w*Bhyp, it could be caused by low coverage when we mapped Illumina short reads to the assembly version 0.2. To our knowledge, there is no record of female-only broods in this species, and sex-ratio distortion was not mentioned when the species was reared under laboratory conditions (DeVries et al., 1999). Here, the specimens used for sequencing were both infected with the same strains and were both males. This could suggest that the *Wolbachia*-induced male-killing phenotype is not expressed or potentially repressed in *B. hypochlora* (Charlat, Reuter, et al., 2007), as was previously described in *Hypolimnas bolina* (Charlat et al., 2005; Charlat, Hornett, et al., 2007). The effect of this *Wolbachia* strain on the reproductive system of this butterfly host and their ecology and evolutionary biology, thus deserves to be further experimentally tested (Charlat, Reuter, et al., 2007; Hiroki et al., 2002; Hornett et al., 2009; Jiggins et al., 2000).

## Conclusion

The assembly of the *Batesia hypochlora* genome assigned 391.297 Mbp (98.87% of the genome) into 15 chromosome-size scaffolds, with an N50 of 25.15 Mbp. The genome is highly complete, with 98.8% of BUSCO represented. We also present 15.54 kbp of mitochondrial assembly and 1.34 Mbp of *Wolbachia* assembly. With the analysis of TE content, gene annotation, and symbiont infection, this genome assembly is a valuable resource for Lepidoptera genomics, ecology, and evolution.

## Ethics approval and consent to participate

Not applicable

## Consent for publication

Not applicable

## Availability of data and materials

Scripts with the commands used are available on github.com/tanpham15/B_hypochlora.

Raw HiFi, Isoseq, and Illumina reads were deposited at ENA (BioProject PRJEB87368). The final nuclear assembly (version 1.0), mitochondrial genome, and annotation are available at NCBI (PRJNA1240833).

## Acknowledgments

We thank Jared Shorma, other field station staff, and volunteers for their help with butterfly collections at Manu Learning Centre, and Carl Yung for DNA isolation advice. We acknowledge NEOF staff and facilities at the University of Sheffield and the University of Liverpool (Centre for Genomic Research). We acknowledge the support of the Freiburg Galaxy Team for the Galaxy server (https://usegalaxy.eu). This research utilised Queen Mary’s Apocrita HPC facility, supported by QMUL Research-IT. http://doi.org/10.5281/zenodo.438045. We acknowledge the assistance of the ITS Research team at Queen Mary University of London. The authors thank Peru’s Servicio Nacional Forestal y de Fauna Silvestre (SERFOR) for permission to conduct field and laboratory research (permit no. D000443-2021-MIDAGRI-SERFOR-DGGSPFFS).

